# The polymorphism of *Hydra* microsatellite sequences provides strain-specific signatures

**DOI:** 10.1101/2020.03.04.977470

**Authors:** Quentin Schenkelaars, Diego Perez-Cortez, Chrystelle Perruchoud, Brigitte Galliot

**Affiliations:** Department of Genetics and Evolution, Institute of Genetics and Genomics in Geneva (iGE3), University of Geneva, Geneva, Switzerland

**Keywords:** *Hydra vulgaris* strains, macerate extract, microsatellite polymorphism, PCR barcoding, cryptic species, speciation

## Abstract

*Hydra* are freshwater polyps widely studied for their amazing regenerative capacity, adult stem cell populations, low senescence and value as ecotoxicological marker. Many wild-type strains of *H. vulgaris* have been collected worldwide and maintained effectively under laboratory conditions by asexual reproduction, while stable transgenic lines have been continuously produced since 2006. Efforts are now needed to ensure the genetic characterization of all these strains, which despite similar morphologies, show significant variability in their response to gene expression silencing procedures, pharmacological treatments or environmental conditions. Here, we established a rapid and reliable procedure at the single polyp level to produce via PCR amplification of three distinct microsatellite sequences molecular signatures that clearly distinguish between *Hydra* strains and species. The TG-rich region of an uncharacterized gene (*ms-c25145*) helps to distinguish between Eurasian *H. vulgaris* strains (*Hm-105*, *Basel1*, *Basel2* and *reg-16*), between Eurasian and North American *H. vulgaris* strains (*H. carnea, AEP*), and between the *H. vulgaris* and *H. oligactis* species. The AT-rich microsatellite sequences located in the *AIP* gene (*Aryl Hydrocarbon Receptor Interaction Protein, ms-AIP*) also differ between Eurasian and North American *H. vulgaris* strains. Finally, the AT-rich microsatellite located in the *Myb-Like cyclin D-binding transcription factor1* gene (*ms-DMTF1*) gene helps to distinguish certain transgenic *AEP* lines. This study shows that the analysis of microsatellite sequences provides a barcoding tool that is sensitive and robust for the identification of *Hydra* strains. It is also capable of identifying cryptic species by tracing microevolutionary events within the genus *Hydra*.

## INTRODUCTION

Since the initial discovery of *Hydra* regeneration by Abraham Trembley in 1744 (1), the freshwater *Hydra* polyp is used as a fruitful model system not only in cell and developmental biology but also for aging, neurobiology, immunology, evolutionary biology and ecotoxicology studies (2–8). *Hydra*, which belongs to Cnidaria, the sister phylum of bilaterians (**Fig 1A**), is closely related to jellyfish although displaying a life cycle restricted to the polyp stage (**Fig 1B**). Over the past 100 years, numerous strains were captured all over the world to explore the variability of the *Hydra* genus and the genetic basis of developmental mechanisms (9–11).

**Fig 1.**
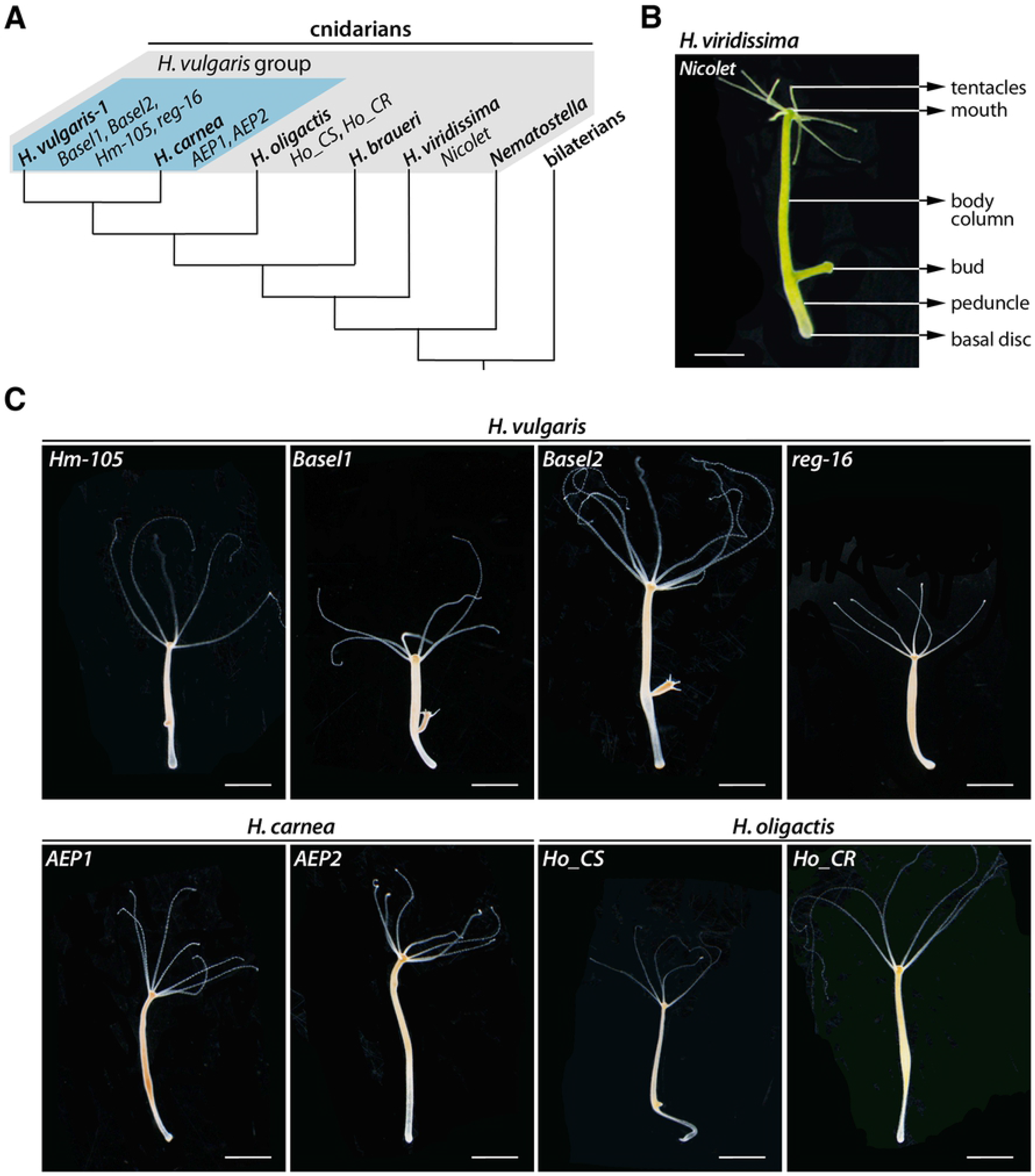
Phylogenetic position and morphology of the freshwater Hydra polyp. **(A)** Phylogenetic tree showing the four *Hydra* species among the *Hydra* genus: *H. vulgaris* and *H. carnea* (blue background), *H. oligactis, H. braueri*, and *H. viridissima*. **(B)** Anatomy of *Hydra*, here a *H. viridissima* polyp from the *Nicolet* strain. *Hydra* polyps exhibit a 0.5-2 cm long tubular structure terminated by the basal disc at the aboral pole and the head at the oral pole. The head region includes a dome structure called the hypostome, terminated by the mouth opening at the tip and surrounded by tentacles at its base. In the lower part of the body column, below the budding zone, the peduncle region precedes the basal disc. Scale bar: 1 mm. **(C)** Morphologies of the *H. vulgaris, H. carnea* and *H. oligactis* strains. Note the presence of a stalk peduncle in *H. oligactis* strains. Scale bars: 2 mm.

The analysis of morphological and cellular criteria identified in *Hydra* strains collected worldwide established four distinct groups named *H. oligactis* (stalked *Hydra*), *H. vulgaris* (common *Hydra*), *H. viridissima* (symbiotic green *Hydra*) and *H. braueri* (gracile *Hydra*) (11) (**Fig 1B, 1C**). The main cellular criterion was provided by the morphology of nematocysts (the venom capsules located inside the mature stinging cells named nematocytes or cnidocytes) that varies between the *Hydra* groups (12). More recently, a series of mitochondrial and nuclear molecular markers were used for barcoding analysis (13–16), which confirmed the relevance of these four groups but also revealed that each group may actually contain several species, e.g. *H. carnea* and formal *H. vulgaris*, also called *H. vulgaris-1*, within the *H. vulgaris* group (**Fig 1A**).

Among the formal *H. vulgaris* species, the *H. magnipapillata* strain 105 (*Hm-105*) is a Japanese strain described by Ito in 1947 (17) and widely used since then (9,14). Several European *H. vulgaris* strains (*Basel, Zürich*, etc.) were also characterized (12), actually found closely related to the Asian *Hm-105* strain. The *AEP* strain, which constitutively produces gametes, was obtained by crossing two North American strains, most likely the *H. carnea* and *H. littoralis* strains, both members of the *H. carnea* species (18,16), subsequently selected for transgenesis (19). Nowadays, the laboratories that use *Hydra* as an experimental model maintain clonal cultures of *H. vulgaris* (*Hm-105*, *Basel, Zürich, reg-16* strains), but also from *H. carnea* (*AEP* strains), *H. viridissima* (e.g. *Nicolet* as Geneva strain) or *H. oligactis* species (*Ho_CS*, *Ho_CR* as European strains) (**Fig 1B, 1C**). A facility located in Mishima (Japan) maintains for the scientific community specimens from a large variety of strains and species (molevo.sakura.ne.jp/Hydra/magni.html).

The importance of identifying the various *Hydra* strains/species relies on the fact that they can exhibit (i) different developmental behaviors, especially the morphogenetic variants that show distinct budding rate or size features in homeostatic context (20–23), (ii) lower regeneration potential such as the *reg-16* strain (24), (iii) abnormal apical patterning such as multiheaded strains (25,26), (iv) specific cellular properties such as the *nf-1* strain that contains neither interstitial stem cells nor interstitial derivatives (27) or the *sf-1* thermo-sensitive strain that loses its cycling interstitial cells upon transient heat-shock exposure (28). Importantly, strains that do not show obvious differences at the morphological or cellular levels actually exhibit variable responses to gene silencing upon RNA interference (29), to drug treatment (30–32) or to environmental stresses (32). In addition, experimental evidences indicate that strain-specific signals regulate the proliferation of interstitial cells (33).

During the past ten years, efforts were made to obtain the *H. vulgaris* genome (34), reference transcriptomes and proteomes (35–37), quantitative RNA-seq in homeostatic and regenerative conditions (38–41) and single-cell transcriptomes (42). Two strains of *H. oligactis*, one undergoing aging (*Ho_CS*) and the other not (*Ho_CR*) were used for transcriptomic and proteomic analysis (32), while genomic sequences were made available for the *H. oligactis* and *H. viridissima* species (41) (See **Table-1**). The current molecular barcoding in *Hydra* is precise and efficient but time-consuming and relatively costly as based on DNA extractions, PCRs amplification followed by DNA sequencing, therefore not well-adapted to large-scale characterization of individual polyps.

**Table-1:**
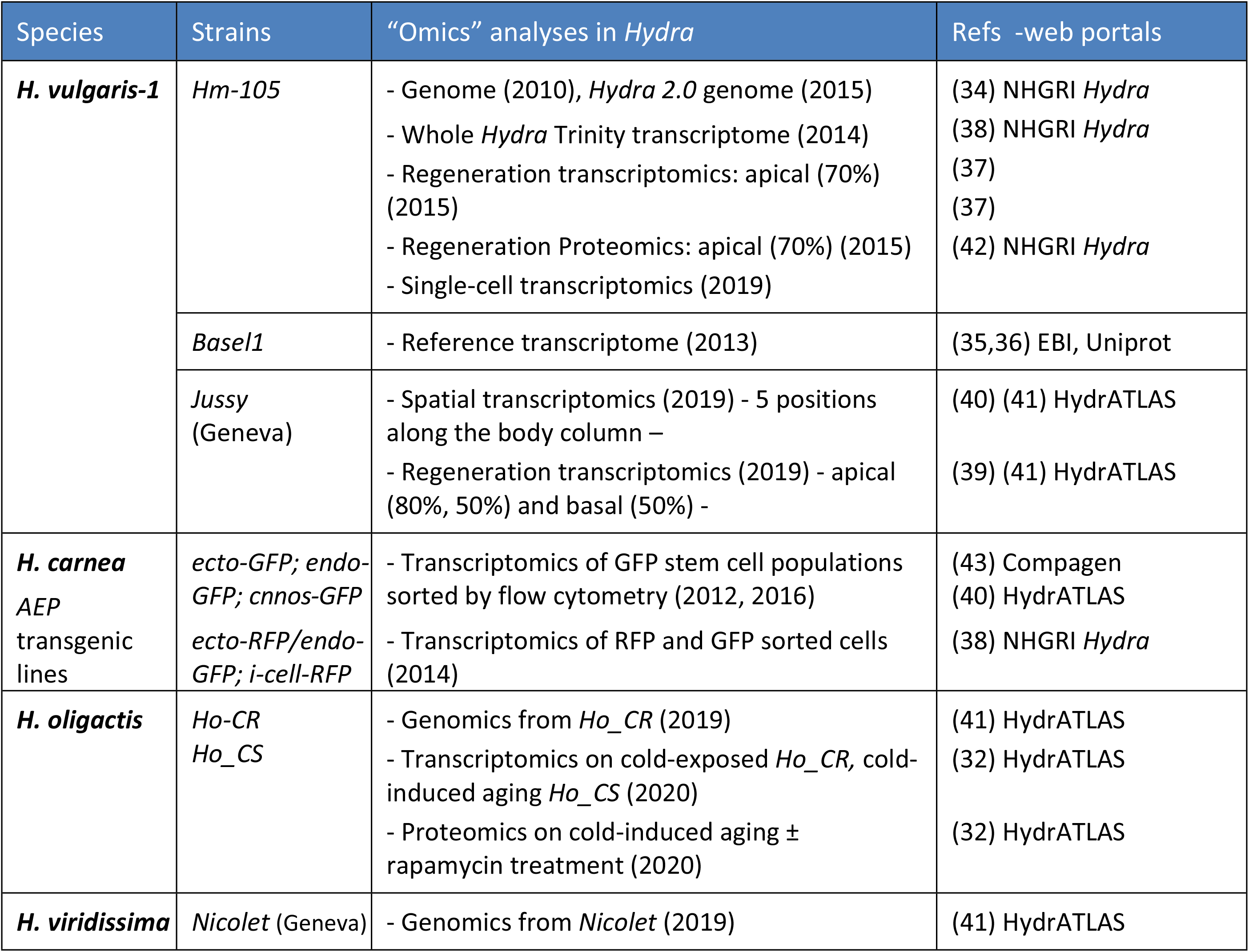
Omics resources in Hydra.

| Species | Strains | “Omics” analyses in *Hydra* | Refs -web portals |
| --- | --- | --- | --- |
| ***H. vulgaris-1*** | *Hm-105* | - Genome (2010), *Hydra 2.0* genome (2015)<br>- Whole *Hydra* Trinity transcriptome (2014)<br>- Regeneration transcriptomics: apical (70%) (2015)<br>- Regeneration Proteomics: apical (70%) (2015)<br>- Single-cell transcriptomics (2019) | (34) NHGRI *Hydra*<br>(38) NHGRI *Hydra*<br>(37)<br>(37)<br>(42) NHGRI *Hydra* |
|  | *Basel1* | - Reference transcriptome (2013) | (35,36) EBI, Uniprot |
|  | *Jussy* (Geneva) | - Spatial transcriptomics (2019) - 5 positions along the body column –<br>- Regeneration transcriptomics (2019) - apical (80%, 50%) and basal (50%) - | (40) (41) HydrATLAS<br>(39) (41) HydrATLAS |
| ***H. carnea* AEP transgenic lines** | *ecto-GFP; endo-GFP; cnnos-GFP* | - Transcriptomics of GFP stem cell populations sorted by flow cytometry (2012, 2016) | (43) Compagen<br>(40) HydrATLAS |
|  | *ecto-RFP/endo-GFP; i-cell-RFP* | - Transcriptomics of RFP and GFP sorted cells (2014) | (38) NHGRI *Hydra* |
| ***H. oligactis*** | *Ho-CR*<br>*Ho_CS* | - Genomics from *Ho_CR* (2019)<br>- Transcriptomics on cold-exposed *Ho_CR*, cold-induced aging *Ho_CS* (2020)<br>- Proteomics on cold-induced aging ± rapamycin treatment (2020) | (41) HydrATLAS<br>(32) HydrATLAS<br>(32) HydrATLAS |
| ***H. viridissima*** | *Nicolet* (Geneva) | - Genomics from *Nicolet* (2019) | (41) HydrATLAS |

See **Table-S3** for details and URLs to access the Compagen, EBI, HydrATLAS, NHGRI *Hydra* web portals.

Microsatellites consist in tandem repeats of short nucleotide motifs of variable length, e.g. (TA)n, (CA/TG)n, (CG)n, (CAG)n, where n represent the number of repetitions (44). These microsatellites are distributed at different locations in the genome, and the number of repeats within a given microsatellite may differ between animals of the same species or population. As a result, microsatellites are widely used for DNA profiling in population genetics studies, but also in criminal investigations, paternity testing, or identification of individuals in the event of a mass disaster (45,46). In these studies, individuals with the same number of repeats at a given genomic location are considered to be closely related, while each additional repeat reflects a divergent step. The combined analysis of different microsatellites makes it possible to construct a genotypic fingerprint specific to each individual, which provides accurate information for tracing evolutionary events such as population bottleneck, migrations, expansions, etc.

The objective of this work was to establish a rapid, inexpensive and reliable method to characterize animals of each strain used in the laboratory. To this end, we established a method that relies on PCR amplification of microsatellite sequences on a single polyp without DNA extraction or sequencing. We show that the analysis of microsatellite polymorphism in animals from either various wild-type strains or transgenic lines provides specific signatures that reliably distinguish strains of the H. vulgaris group. This barcoding method, now routinely applied in our laboratory, is efficient and well suited for large-scale studies.

## Materials and Methods

### Hydra strain collection

The wild-type strains used in this study were a kind gift from colleagues, *Basel1* and *AEP1* from B. Hobmayer (University of Innsbruck), *Basel2*, *Hm-105* and *Ho_CR* from T. Holstein (University of Heidelberg), *AEP2* from R. Steele (University of California), *Ho_CS* from H. Shimizu (National Institute of Genetics, Mishima) and *Nicolet* from Mr. Nicolet (Geneva). The AEP transgenic lines that constitutively express GFP in their epithelial cells, either gastrodermal (*endo-GFP*) or epidermal (*ecto-GFP*), were produced by the Bosch Lab (University of Kiel) (19,47) (Wittlieb et al., 2006, Anton-Erxleben et al., 2009) and kindly provided to us. The *AEP1* transgenic lines expressing the *HyWnt3**–**2149::GFP* construct (here named *Wnt3::GFP*) either in epidermal or gastrodermal epithelial cells were produced in-house with the *HyWnt3**–**2149::GFP-HyAct:dsRed* reporter construct kindly given by T. Holstein (48,49). We also produced in the AEP2 strain the Q82-203 and Q82-293 lines by injecting early embryos with the *HyActin:Q82-eGFP* construct (QS, unpublished) following the original procedure (19). All cultures were fed three times a week with freshly hatched *Artemia* and washed with *Hydra* Medium (HM) (24) (Sugiyama and Fujisawa, 1977a).

### One-step preparation of macerate extracts

Polyps were washed three times five minutes in distilled water. Then, single polyps were dissociated into 50 μL distilled water by energetically pipetting them up and down until there is no tissue left, and immediately transferred on ice. Cell density of each macerate was estimated by measuring the OD_600_ using a NanoDrop One (Thermo Sientific). The DNA content and DNA purity were roughly estimated by measuring the absorbance of each sample at 230, 260 and 280 nm. To implement an efficient one-step PCR procedure, we selected three *AEP2* polyps showing a regular size (about 4-6 mm long without the tentacles).

### PCR amplification from macerate extracts

To test the efficiency of PCR amplification on macerate extracts, we used primers of the *β- actin* gene (**Table-S1**) on 0, 0.5, 1.5, 5 and 15 μL macerate extract as template for a final 25 μL PCR mix (1x Taq Buffer, 1x Coral Load, 400 nM of each primer, 160 nM dNTPs and 0.5 unit of Top Taq Polymerase, Qiagen). Subsequently we used 5 μL out of 50 μL macerate extract to amplify the mitochondrial *cytochrome C oxidase I* (*COI*) gene, the mitochondrial *16S ribosomal DNA* (*16S*) and the microsatellite regions (ms) in each strain (**Table-S1**). After an initial denaturation step at 94°C for two minutes, samples were submitted to 30 cycles of (i) denaturation at 94°C for 15 seconds, (ii) annealing at 52°C for 30 seconds and (iii) a 30-60 seconds elongation step at 72°C. The process was terminated by a final extension at 72°C for 15 minutes. 10 μL PCR products were run on a 2.5% agarose gel at 120 V for two to three hours in the case of microsatellites, stained with ethidium bromide and revealed under UV-light.

### Cloning and sequencing

For sequence validation, the PCR products were cloned using the pGEMT kit (Promega): 3 μl PCR products were ligated to 50 ng pGEMT vector in the presence of 3 units T4 ligase overnight at 18°C (final volume 10 μL). Plasmidic DNA was integrated into competent DH5α *E. coli* and colonies were screened thanks to alpha-complementation. After overnight culture, plasmidic DNA was extracted using the CTAB procedure and sequenced using standard T7 primer at Microsynth (Basel, Switzerland). The number of colonies we sequenced and their origin (single or several animals) is indicated for each microsatellite sequence in **Table-S3**.

### Phylogenetic analyses

The *COI* and *16S* genes were selected for phylogenetic analyses. Corresponding DNA sequences were amplified by direct PCR amplification method as described above and sequenced (**Table-S2**). The obtained sequenced were aligned with the dataset previously produced by Martinez et al. (15) using the ClustalW function of BioEdit v7.2.6.1, and Maximum Likelihood phylogenetic trees were constructed with the PhyML 3.0 software (http://www.atgc-montpellier.fr/phyml/) applying the GTR substitution model (50). The robustness of the nodes was tested by 1000 bootstraps.

## RESULTS

### One-step genomic amplification after quick mechanical tissue maceration

To bypass genomic and mitochondrial DNA extractions that are time-consuming and expensive when massively performed, we established a rapid animal dissociation in water that provides genomic DNA of sufficient quantity and quality for PCR reaction. Although the efficiency of such PCRs certainly varies with the gene of interest, the primers and the size of the amplicon, we obtained PCRs using macerate extracts resulted in strong bands for *β-actin* (193 bp), implying that the application of a mechanical force to dissociate the tissues and the denaturation step at the beginning of the PCR are sufficient to release high quality genomic DNA to allow the amplification of the target sequence (**Fig 2A**). More precisely, despite slight variations in band intensity, certainly reflecting the amount of starting material, the amplification remained highly efficient whatever the polyp and the template volume used here. Accordingly, for all subsequent experiments, we used one tenth of macerate extract as template for *COI*, *16S* and microsatellite amplifications (**Fig 2B**). We also obtained efficient PCR amplification from macerate extracts prepared from fixed animals stored at -20°C for several years, especially for mitochondrial DNA amplification. This procedure thus allows us to gain genetic information from fresh as well as old samples.

**Fig 2.**
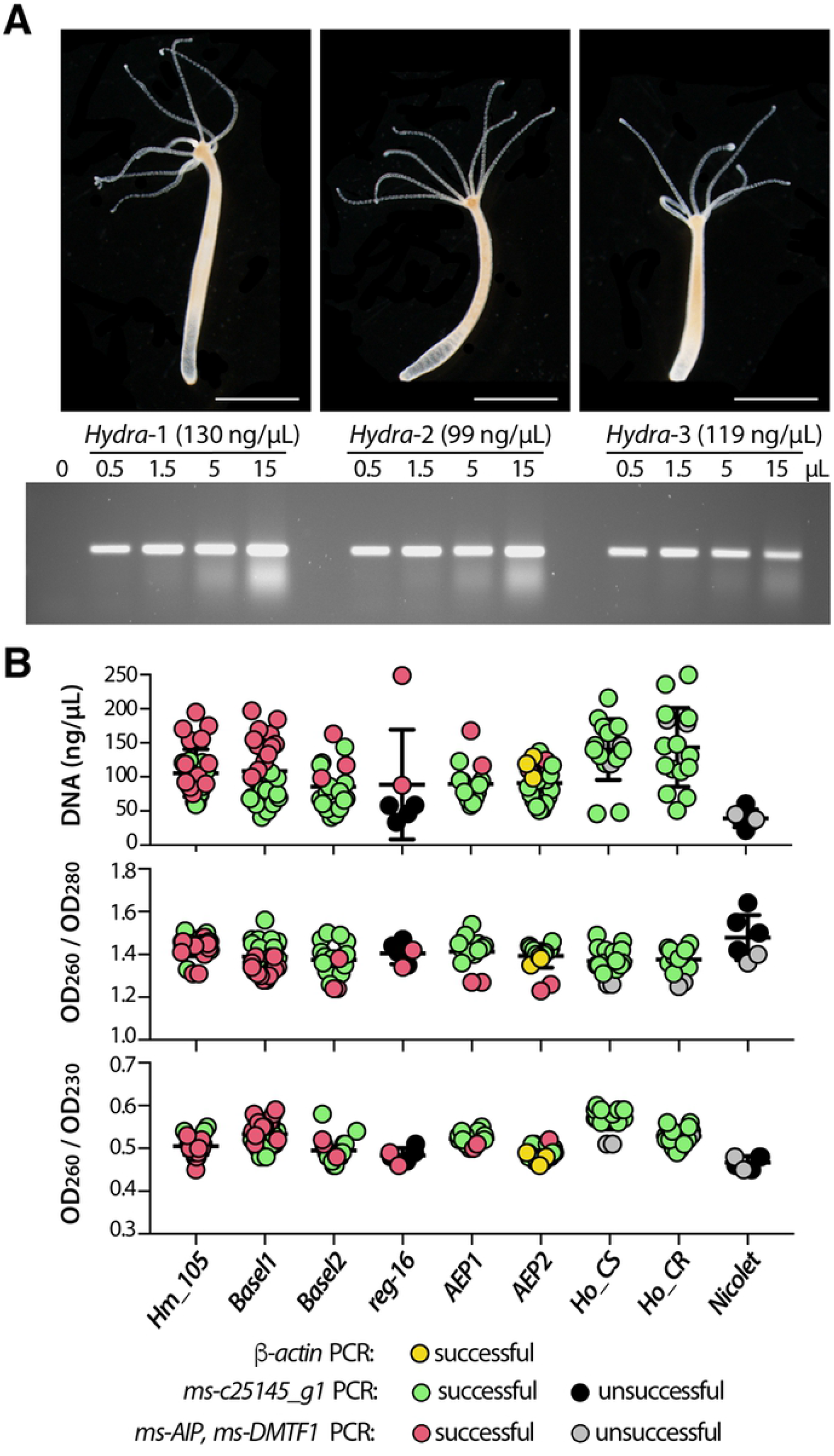
Direct genomic DNA amplification from single Hydra polyp. **(A)** Efficacy of PCR amplification of *ß-actin* genomic DNA according to the original size of the dissociated polyp. DNA from each polyp was resuspended in 50 μl. Scale: 2 mm. **(B)** Graphic representation of DNA extraction efficiency and DNA purity as deduced from OD measurements at 260, 230 and 280 nm wave lengths. Each dot represents a value obtained from a single polyp. For each DNA, the efficiency of PCR amplification is indicated with a color code.

### Phylogenetic assignation of Hydra strains to the different groups and species

Next, we confirmed the assignation of each strain we acquired to one of the four *Hydra* groups previously described (i.e. *H. vulgaris, H. oligactis*, *H. viridissima*, *H. braueri*), and when relevant to the species identified within each group, namely *H. vulgaris 1* and *H. carnea* within the *H. vulgaris* group (13–16). Briefly, we performed phylogenetic analyses of the *COI* and *16S* sequences, efficiently amplified from one single polyp per strain of interest (*AEP1, AEP2, Basel1, Basel2, Hm-105, reg-16, Ho-CR, Ho-CS, Nicolet*) as detailed above. The global topology of the *COI* tree retrieves the four orthologous groups (**Fig 3**), which is not the case in the *16S* analysis where the *H. vulgaris* group actually includes the *H. brauerei* and *H. oligactis* groups that thus do not appear monophyletic (**Fig S1**).

**Fig 3:**
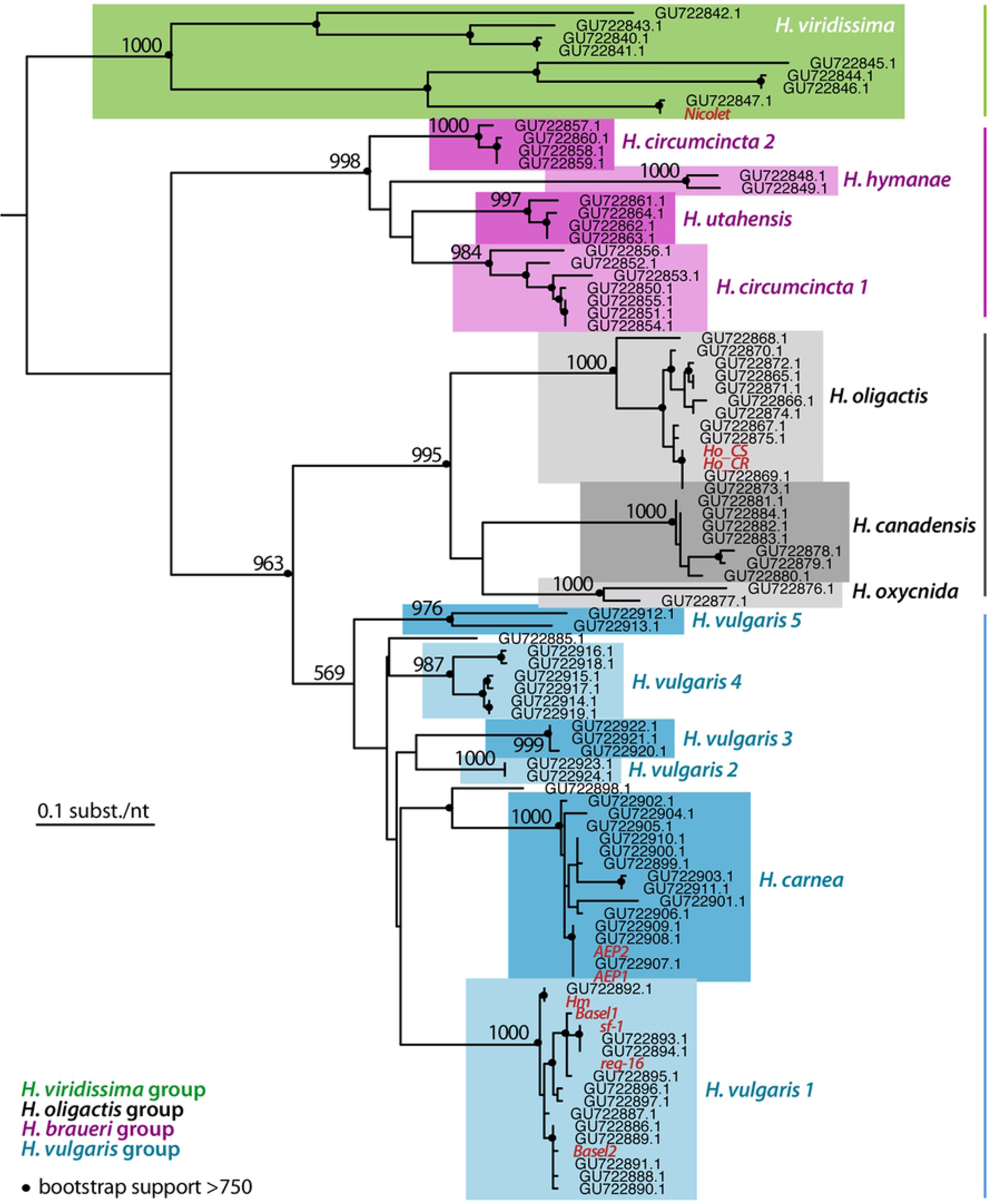
Phylogenetic relationships within the Hydra genus based on the analysis of the Cytochrome Oxydase I (COI) DNA sequences. The maximum likelihood tree of the *COI* sequences was built by adding to the dataset of 85 COI sequences available on Genbank (15) the 10 sequences obtained in the present study (written in red, see **Table-S3** for accession numbers). Black dots indicate the robustness of the nodes as deduced from the bootstrap support (at least 750 over 1’000 bootstraps). This tree confirms the presence of four distinct *Hydra* groups (*H. viridissima, H. braueri*, *H. oligactis and H. vulgaris*). Within the *H. vulgaris* group, note the position of the *AEP* sequences within the *H. carnea* sub-group.

However, in both analyses, the sequences of the strains tested were grouped as expected within the 13 species previously identified (i.e. *H. circumcincta 1 and 2, H. hymanae, H. utahensis, H. oligactis, H. canadensis, H. oxycnida, H. carnea* and *H. vulgaris* 1 to 5). The *Hm- 105* and *Hv_Basel* sequences are grouped in the *H. vulgaris 1* group, a Eurasian group which contains the *Hm-105* reference sequences (GU722892.1 for COI and GU722807.1 for 16S), the *H_CS* and *Ho_CR* sequences both belong to the *H. oligactis* group, and the *Nicolet* sequences belong to the *H. viridissima* group. This analysis also confirms that the *AEP* sequences (*AEP1, AEP2*) belong to the *H. carnea* group that contains all the sequences of the North American strains of the *H. vulgaris* group and only these sequences (**Fig 3**). We found that the genomic *16S* sequences of the two *Hv_Basel* strains are identical, while the mitochondrial *COI* sequences are different with nine out of 657 bp mismatches (sequences obtained twice independently). Consequently, animals of these two cultures can be considered as belonging to two different strains, which we have named *Basel1* and *Basel2*. In contrast, the *COI* and *16S* sequences of *AEP1* and *AEP2* were identical, suggesting that they could represent a single strain.

### Identification of three microsatellite regions in the Hydra genome

We then analyzed some microsatellite sequences to test the conclusions obtained in the phylogenetic analyses and to establish a method for easy identification of strains belonging to the *H. vulgaris* group. To identify *H. vulgaris* genomic regions that contain microsatellites, we blasted two different tandem repeat motifs (TA)_15_ and (CA)_15_ against *AEP* transcriptomes available at the HydrATLAS web portal. We found three transcripts expressed by *AEP* polyps that encode repeats, the first one *c25145_g1_i04* contains TG-repeats in its first intron (**Fig 4, Fig S2**), the second *c8134_g1_i1* encodes the Aryl-hydrocarbon receptor-Interacting Protein (AIP) and contains AT-repeats in its 5’ untranslated region (UTR) (**Fig 5, Fig S5**), and the third one (*c21737_g1_i4*) encodes the cyclin-D-binding Myb-like Transcription Factor 1 (DMTF1) and contains AT-repeats located in the 3’UTR (**Fig 6, Fig S6**).

**Fig 4:**
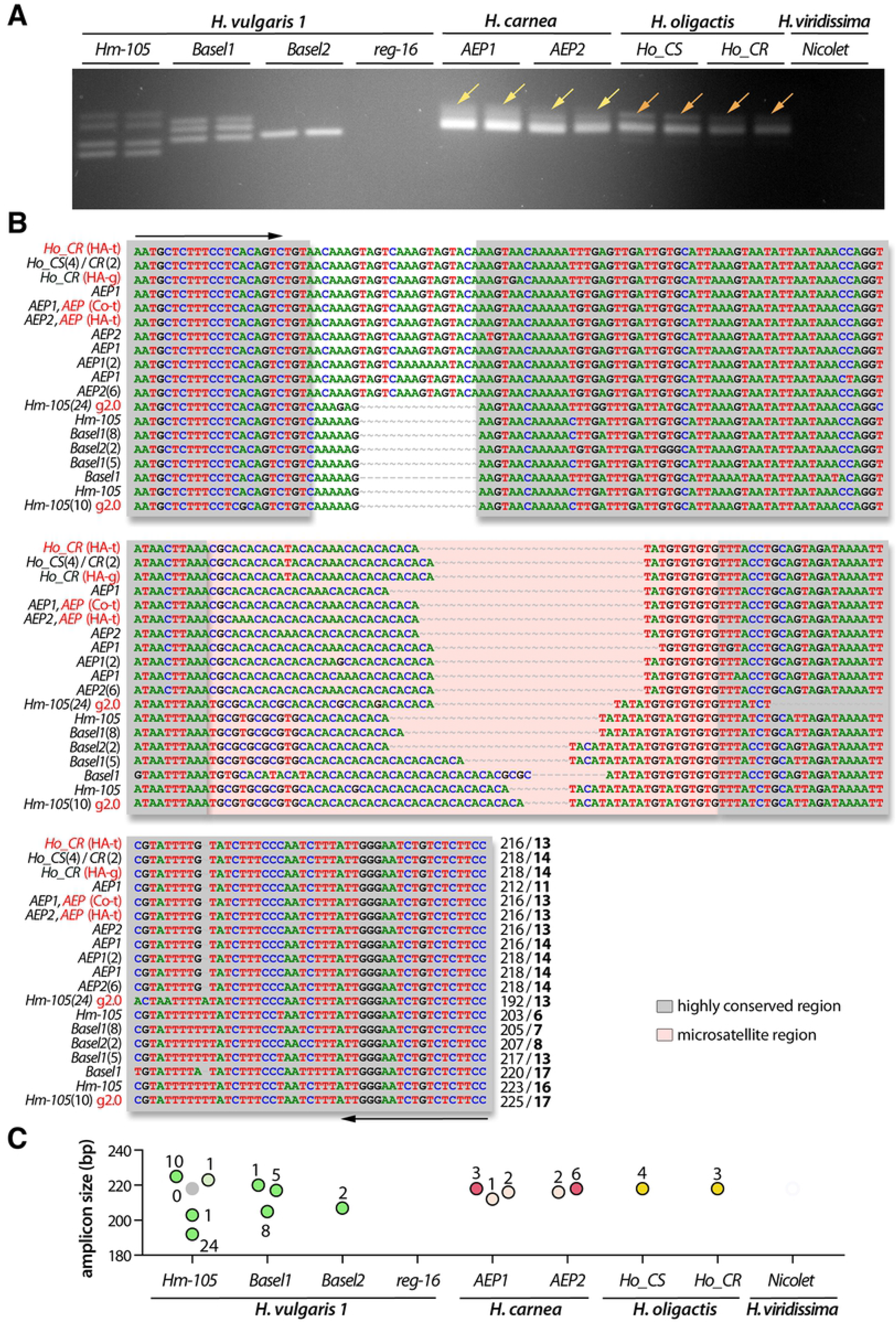
Analysis of the polymorphism of the TG-rich microsatellite c25145 sequence (ms-c25145) **(A)** Amplification of the *ms-c25145* genomic sequences from seven out of nine tested strains that represent three *Hydra* sub-groups, *H. vulgaris1* (*Hm-105, Basel1, Basel2* strains), *H. carnea* (*AEP1*, *AEP2* strains) and *H. oligactis* (*Ho_CR, Ho_CS* strains). Yellow arrows point to a smear detected in both *AEP1* and *AEP2*, orange arrows point to a faint second band detected in both *Ho_CS* and *Ho_CR*. **(B)** Sequence alignment of the *ms-c25145* region. The salmon-pink color box indicates the central TG-rich microsatellite region embedded within highly conserved regions (grey boxes). Primer sequences used for amplification are indicated with black arrows. Numbers in brackets after the strain name indicate the number of independent positive sequencings, numbers at the 3’ end indicate the size of the PCR product and the number of TG-repeats (bold). Red writings indicate transcriptomic (t) or genomic (g) sequences available on the HydrATLAS (HA) server, NHGRI *Hydra* web portal for the *Hydra* 2.0 genome (g2.0), or Compagen (Co) server (see **Table-S3**). **(C)** Graphical representation of the different *ms-c25145* amplicons as deduced from sequencing data. Each dot corresponds to a distinct amplicon confirmed by one or several sequencings as indicated by the number of sequenced colonies (see **Table-S3**). Green, red and yellow color dots correspond to expected sizes, lighter color dots refer to sequences with errors (PCR or sequencing), the grey dot indicates missing data.

**Fig 5:**
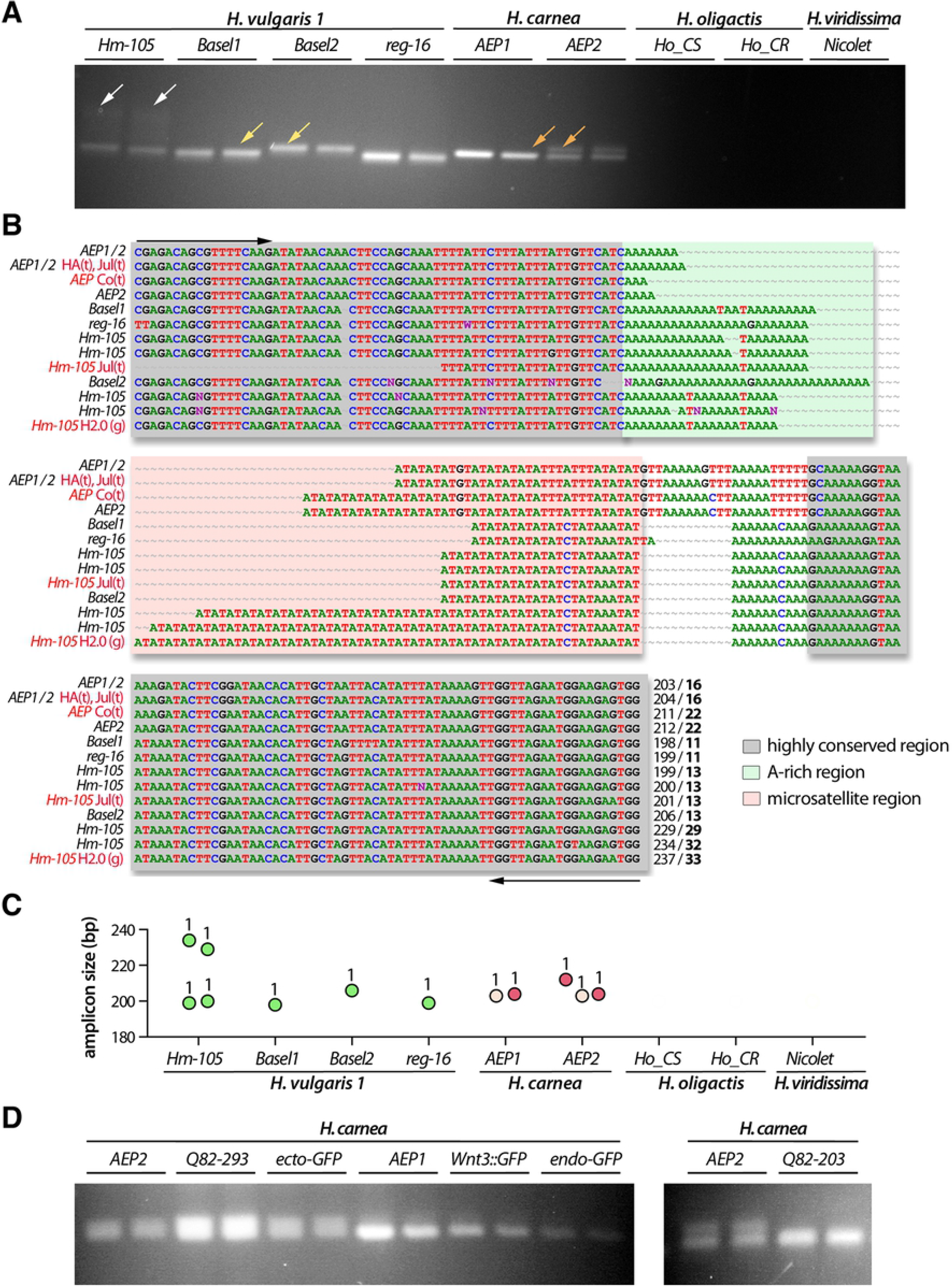
Analysis of the polymorphism of the AT-rich microsatellite region of the Aryl-hydrocarbon receptor-Interacting Protein gene (ms-AIP) **(A)** Amplification of the *ms-AIP* genomic sequences in six out of nine tested strains, which represent two distinct *H. vulgaris* sub-groups, *H. vulgaris 1* (*Basel1, Basel2, Hm-105, reg-16*) and *H. carnea* (*AEP1, AEP2*). White arrows point to a faint band observed only in *Hm-105* polyps, yellow arrows indicate a size difference between *Basel1* and *Basel2*, and the orange arrows show a second band detected in *AEP2* but not in *AEP1*. **(B)** Alignment of the *ms-AIP* sequences. The color boxes indicate the AT-rich central region (salmon-pink) and an A-rich motif (green) embedded within highly conserved regions (grey). Primer sequences used for amplification are indicated with black arrows. Numbers at the C-terminus indicate the PCR product size and the number of AT-repeats (bold). Red writings indicate transcriptomic (t) or genomic (g) sequences available on HydrATLAS (HA), NHGRI web portal for the *Hydra* 2.0 genome (g2.0) and Juliano transcriptomes (Jul), or Compagen (Co) server (see **Table-S3**). **(C)** Graphical representation of the *ms-AIP* amplicons as deduced from sequencing data. Dot legend as in **Fig 4**. **(D)** Amplification of *ms-AIP* in five transgenic lines *ecto-GFP* and *endo-GFP* produced in uncharacterized AEP (REFs), *AEP1_Wnt3* (Vogg et al. 2019), *AEP2_203* and *AEP2_293* (QS, unpublished).

**Fig 6:**
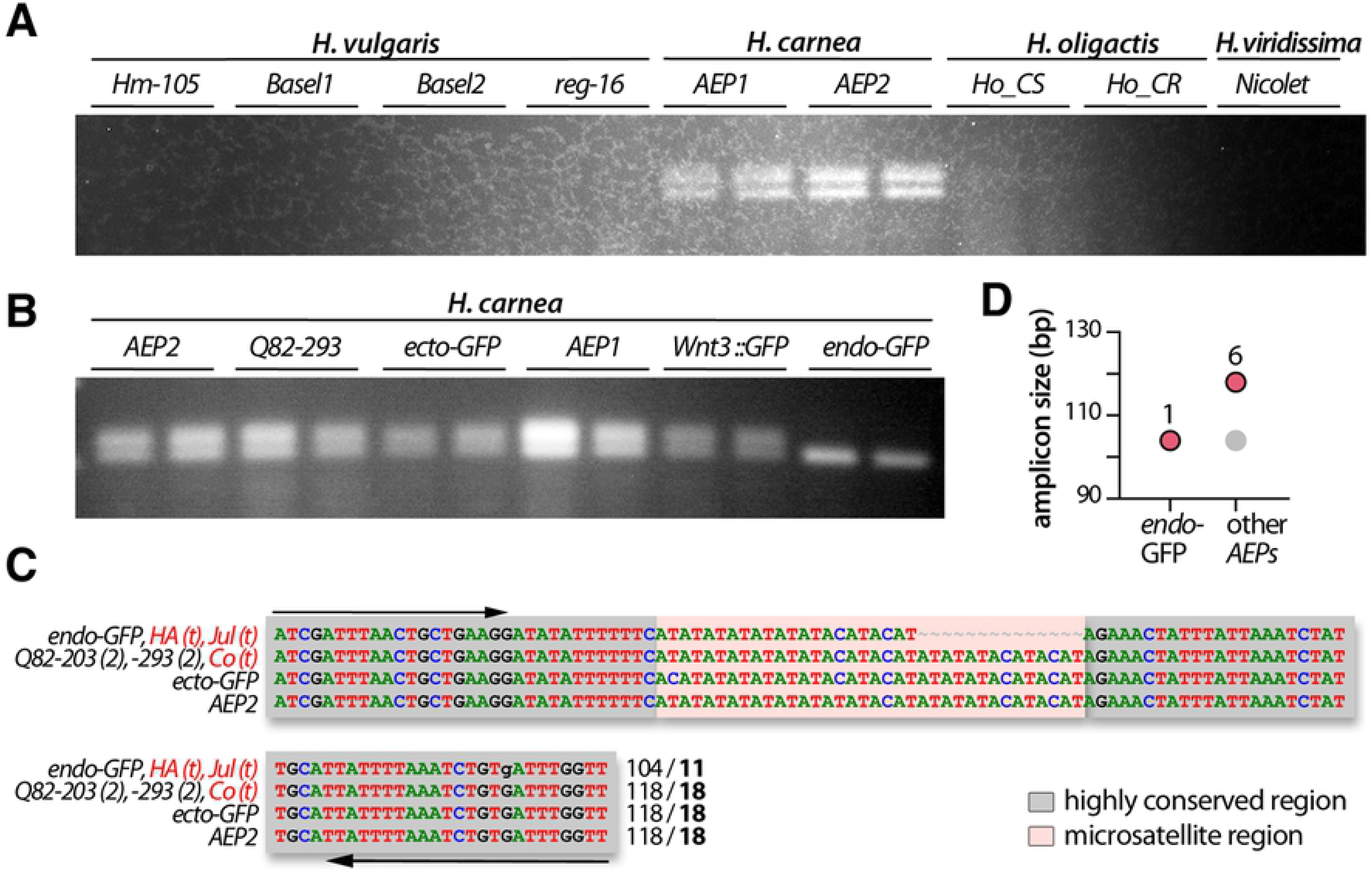
Analysis of the polymorphism of the AT-rich microsatellite detected in the Cyclin-D-Binding Myb-Like Transcription Factor 1 gene (ms-DMTF1) **(A, B)** Amplification of the *ms-DMTF1* genomic sequence is restricted to the *AEP* strains, either unmodified (*AEP1, AEP2*) or transgenic (*Q82-293*, *ecto-GFP*, *Wnt3::GFP*, *endo-GFP*) lines. **(C)** Alignment of the *ms-DMTF1* sequences. The color boxes indicate the AT-rich central region (salmon-pink) embedded within highly conserved regions (grey). Primer sequences used for amplification are indicated with black arrows. Numbers in brackets after the strain name indicate the number of independent positive sequencings, numbers at the 3’ end indicate the size of the PCR product and the number of AT-repeats (bold). Red writings indicate transcriptomic (t) or genomic (g) sequences available on HydrATLAS (HA) server, NHGRI web portal for the *Hydra* 2.0 genome (g2.0) and Juliano transcriptomes (Jul), or Compagen (Co) server (see **Table-S3**). **(D)** Graphical representation of the size of the *ms-DMTF1* amplicons as deduced from sequencing data. Red color dots correspond to expected sizes, the grey dot indicates missing data.

Next, we validated these sequences onto genomic and transcriptomic databases publicly available for *Hm-105* on National Human Genome Research Institute (NHGRI) and Compagen. These three microsatellite regions were selected as they were retrieved from most databases and contained a variable number of microsatellite repeats between *H. vulgaris 1 (Hm-105)* and *H. carnea (AEP)*. We named these microsatellite regions *ms-c25145*, *ms-AIP* and *ms-DMTF1* respectively; access to the corresponding transcriptomic and genomic sequences are given in **Table-S3**.

### The ms-c25145 polymorphism helps to discriminate between Hydra species and H. vulgaris strains

The TG-rich *ms-c25145* could be detected within two different *Hm-105* genomic regions (*Sc4wPfr_1246, Sc4wPfr_396* scaffolds) and the direct PCR approach efficiently amplified the *ms-c25145* genomic sequences in seven strains (*Hm-105, Basel1, Basel2, AEP1, AEP2, Ho_CS, Ho_CR*), but remained inefficient in the *reg-16* strain (*H. vulgaris* group) and the *H. viridissima* strain named *Nicolet*, possibly due to mismatches into primer regions (**Fig 4A, Fig S3**). The patterns obtained for *ms-c25145* are quite different between *Hm-105* (four bands), *Basel1* (three bands) and *Basel2* (single band), indicating that these strains can indeed be considered as distinct, in agreement with the results of the *COI* phylogeny (**Fig 3**). Concerning the *AEP1* and *AEP2* strains, the *ms-c25145* patterns appear quite similar, with a main band about 216 bp long, and a smear of larger and less intense bands (**Fig 4A**, yellow arrows). This pattern is quite distinct from the sharp bands observed in *Basel2*. An intense band of similar size than in *AEPs* (218 bp) is observed for *Ho_CS* and *Ho_CR* as well as some weaker and longer amplicon (**Fig 4A**, orange arrows). In summary, *ms-c25145* appears as an informative marker to distinguish *Hm-105, Basel1* and *Basel2* strains from each other, and from strains representative of the *H. carnea, H. oligactis* and *H. viridissima* species.

To confirm these results, we cloned the PCR products and randomly sequenced some colonies from at least two animals of each strain (for sequencing details see **Table-S3**), and we found the sequence size fully consistent with the observed size of the bands on the gels (**Fig 4B, 4C**). Indeed, the lowest *Basel1* PCR product is slightly shorter (205 bp) than the unique *Basel2* PCR product (207 bp), whereas the two other *Basel1* PCR products are 217 and 220 bp long. For *Hm-105*, we retrieved sequences for three PCR products out of the four observed on the gel, two corresponding to the shortest bands (192 and 203 bp) and one to the upper one (223-225 bp). In *AEP* samples, we retrieved multiples sequences with nucleotide polymorphism (AA instead of CA repeat) that correspond to the most abundant PCR product, ranging from 212 to 218 pb. Finally, sequencing results confirmed that the main PCR product observed in *H. oligactis* strains correspond to the 218 bp band, also found in *AEPs*. The sequencing data provided robust results regarding the number of TG-repeats of each sequence, i.e. 6, 13 and 17 in *Hm-105*, 7 and 13 in *Basel1*, 8 in *Basel2*, and 14 in the *H. carnea* and *H. oligactis* sequences.

We also analyzed the location of this *ms-cv25145* microsatellite sequence within the *ms-cv25145* gene: It appears intronic, located after the first exon, about 245 bp downstream to the 5’ end (**Fig S2**). The *c25145* gene encodes a putative evolutionarily-conserved protein with an unknown function as deduced from the alignment of the *Hydra* c25145 deduced protein product with related bilaterian sequences (**Fig S4**). We found similarities in the N-terminal moiety (~100 first amino acids) with hypothetical proteins expressed by the sea cucumber *Apostichopus japonicus* (51), the arthropods *Folsomia candida* and *Sipha flava* (aphid), the mollusc *Crassostrea gigas*, the teleost fish *Myripristis murdjan, Sinocyclocheilus rhinocerous* or *Danio rerio*. Within this domain, a signature can be identified, formed of 37 residues, from which 32 are present in the *Hydra* protein (**Fig S4**).

### The ms-AIP polymorphism helps to identify H. vulgaris and H. carnea strains

The second microsatellite region (*ms-AIP*) is an AT-rich region located in the 5’UTR region of the gene encoding the Aryl-hydrocarbon (AH) receptor-Interacting Protein (**Fig S5**). The polymorphism of *ms-AIP* is more restricted than that of *ms-c25145*, as we were unable to amplify these genomic sequences from the *H. oligactis* and *H. viridissima* strains (**Fig 5A**). Nevertheless, *ms-AIP* is useful to discriminate between the strains within the *H. vulgaris group*, i.e. *Hm-105, reg-16, Basel1, Basel2, AEP1, AEP2*. Two PCR products were obtained after genomic amplification from *Hm-105* and *AEP2* whereas a single PCR product was amplified from the other strains, with a specific size for each strain (**Fig 5A**).

The sequencing results mainly matched with the patterns detected by electrophoresis (**Fig 5B, 5C**), proving that distinct band sizes reflected stable strain-specific variations in both the length of the A-rich region and the number of AT-repeats. Indeed, two distinct batches of sequences were obtained for *Hm-105* (199-200 and 229-234 pb; 13 and 29-33 AT-repeats respectively). The slight differences observed in the amplicon size among a given animal possibly resulted from polymerase slippage during the PCR process or from an altered sequencing process, as often observed in AT-rich regions (**Fig 5B**). In addition, the *ms-AIP* sequences obtained from *Basel1*, *Basel2* and *reg-16* are consistent with the 198, 206 and 199 pb long bands observed on the gels, corresponding to 11, 13 and 11 AT-repeats respectively. In contrast to *ms-c25145*, the analysis of the *ms-AIP* sequences helps distinguish between *AEP1* and *AEP2*, since *AEP2* shows two bands, 204 and 212 bp long corresponding to 16 and 22 AT-repeats, while only the lowest band is present in *AEP1* (**Fig 5A**, orange arrow). As a consequence, we consider *AEP1* and *AEP2* as two distinct strains even though their *COI* and *16S* sequences are identical (**Fig 2**). Since we were able to identify different patterns in the *AEP1* and *AEP2* strains, we also looked at the *ms-AIP* polymorphism in *AEP* transgenic lines (**Fig 5D**). The *Q82-293* and *ecto-GFP* lines show the two-bands pattern found in *AEP2* while the *Wnt3::GFP*, *endo-GFP* and *Q82-203* lines show the same single-band pattern than *AEP1*. In summary, the analysis of the *ms-AIP* patterns are informative to identify and characterize strains of the *H. vulgaris 1* species. In addition, in contrast to *ms-c25145, ms-AIP* provides a useful marker for the *AEP* strains and *AEP* transgenic lines.

### The ms-DMTF1 microsatellite helps to discriminate between the H. carnea AEP lines

The third microsatellite sequence (*ms-DMTF1*) is also AT-rich but located in the 3’ UTR of the *cyclin-D-binding Myb-Like transcription factor 1* gene (**Fig S6**). The *ms-DMTF1* primers were designed for *H. carnea* strains and are thus only suitable for strains that belong to the *H. vulgaris* group (**Fig 6A**). Accordingly, they are useful to discriminate between animals of this *H. vulgaris* group. The analysis of the *ms-DMTF1* polymorphism does not show variability between *AEP1* and *AEP2* but remains useful to distinguish the *endo-GFP* transgenic animals from all other *AEPs* (**Fig 6**). In fact, all the AEP strains and lines we tested here but the transgenic line *endo-GFP*, provide a two-band pattern, the lowest band being similar in size with the single one found in the *endo-GFP* (**Fig 6B**). By shot-gun sequencing of the PCRs products from different *AEP* animals, we found that the sequence of the upper band is 118 bp long (**Fig 6C, 6D**). In complement, the sequencing data obtained in the *endo-GFP* animals identified a PCR product that corresponds to 104 bp. Interestingly, available transcriptomes confirm the existence of both sequences (**Fig 6C** and **Table-S3**).

### Comparative analysis of the information brought by microsatellite barcoding

To establish the respective barcode values of the *ms-c25145, ms-AIP* and *ms-DMTF1* microsatellites (**Fig 7**), we compared the results obtained in the 36 strain/species pairs tested for each microsatellite. From the analysis of these three microsatellites we deduced four levels of information, (1) informative when the patterns are distinct between the two strains/species, (2) partially informative when microsatellite amplification is observed in one strain/species but not in the other, (3) or when the patterns obtained are identical between the two strains/species, (4) non-informative when amplification is not observed in either strain/species.

**Fig 7:**
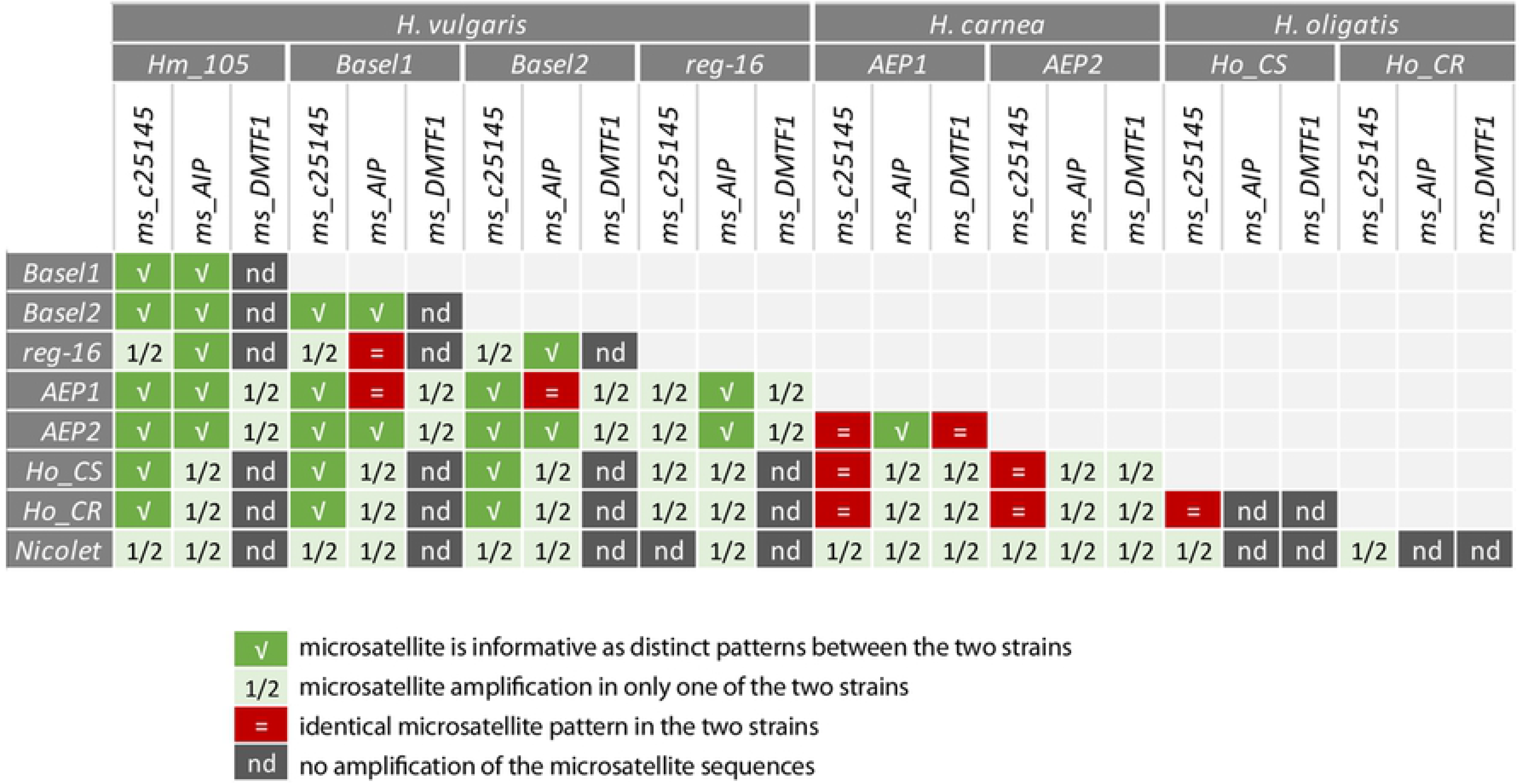
Summary scheme showing the value of each microsatellite for efficient discrimination between Hydra species and Hydra strains

Among these three microsatellites, *ms-c25145* is the most informative as the only one amplified in three distinct species (*H. vulgaris 1, H. carnea, H. oligactis*), providing a positive discrimination in 29 pairs (80.6%), either based on specific patterns as observed in 15 pairs (41.7%) or on an amplification restricted to a single strain/species in 14 pairs (38.9%). The *ms-AIP* is amplified in *H. vulgaris 1* and *H. carnea*, providing a positive discrimination in 30 pairs, based on specific patterns in only 12 pairs (40%) and on an amplification restricted to a single strain/species in 18 pairs (60%). Finally, *ms-DMTF1* is only amplified in the *AEP1* and *AEP2* strains, providing a similar pattern in eight pairs, but a distinct one in some transgenic strains. We concluded that the approach presented here fulfilled our initial objective since it allowed us to properly characterize all strains of the *H. vulgaris* group used in our laboratory, i.e. strains *Hm-105, Basel1, Basel2* and *reg-16* of the species *H. vulgaris-1* as well as strains *AEP1, APE2* of the species *H. carnea*. By contrast, the phylogenetic approaches based on *COI* and *16S* sequences had failed as the *COI* and *16S* sequences were identical between some strains.

### Analysis of speciation events in H. vulgaris based on the microsatellite signatures

Although the region surrounding the microsatellites sequences is quite conserved between all strains, we observed systematic differences between *H. vulgaris 1* and *H. carnea* strains in the organization of the amplified regions such as the TAGTCAAAGTAGTACA deletion in the upstream non-conserved region of *ms-c25145* in *H. vulgaris 1* strains (**Fig 4B**), or the size difference in the A-rich region in *ms-AIP* (**Fig 5B**). The conserved deletions in one of the two subgroups and the differences in the microsatellite motifs suggest that the genetic flux between *H. carnea* strains *(AEP*) and *H. vulgaris 1* strains (*Hm-101*, *Basel1, Basel2, reg-16*) no longer exists, suggesting that *H. vulgaris 1* and *H. carnea* can be considered as two cryptic species (52). This hypothesis requires further confirmation that could be obtained by amplifying the *ms-c25145*, *ms-AIP* and additional microsatellite sequences from representative animals of the 14 hypothetical species reported by Schwentner and Bosch (14).

## DISCUSSION

### The direct dissociation of soft tissues provides quality templates for genomic PCR amplification

Genomic extractions for multiple samples as well as for population genetics studies can be rapid but costly when commercial kits are used, or time-consuming and risky when reagents that are rather toxic to humans and/or the environment are used (e.g. guanidium thiocyanate, ß-mercaptoethanol). For these reasons, we have tried here to bypass the genomic extraction step and to use directly as PCR substrate dissociated *Hydra* tissues that we call “macerate extracts”. The rapid and inexpensive protocol we present here is based on the mechanical dissociation of the tissues, which reliably allows the PCR amplification of mitochondrial and nuclear DNA. This procedure is now commonly used in our laboratory, not only to amplify microsatellite sequences and detect in *Hydra* cultures suspected contamination by polyps of other strains, but also to amplify genomic sequences of genes of interest for directly sequencing or insertion into plasmid vectors. We also successfully applied this procedure to fixed *Hydra* tissues as reported above, as well as poriferan larvae (e.g. *Oscarella lobularis*, not shown). Therefore, this protocol can be effectively applied to soft tissues from any developing or adult organisms, especially when small amounts of tissue are available.

### Systematize characterization of Hydra strains to improve data reproducibility

The microsatellite barcoding approach reported here offers a series of important advantages in that it is (i) *sensitive*, detecting a 2 bp shift in amplicon size, (ii) *simple*, requiring no chemicals or materials other than those used in ordinary PCR as in conventional barcoding approaches, (iii) *fast*, with data being acquired in less than a day, (iv) *robust* as it provides reproducible results. The immediate use of macerate extracts could be a possible limitation of this procedure. Indeed, we did not test the quality of these macerate extracts after their storage in a frozen state, assuming that nucleic acid degradation would occur. Nevertheless, we were able to amplify genomic DNA obtained after mechanical dissociation from intact frozen animal samples, implying that fresh material is not an absolute requirement.

In the context of life sciences where reproducibility can be a challenge (53,54), the development of tools to properly characterize the animals we work with appears to be a cornerstone towards more effective research. Indeed, *Hydra* laboratories use a wide variety of strains that are known to respond differently to chemical treatments or show variable sensitivity to gene expression silencing by RNAi. This procedure opens up the possibility of conducting blind clonal culture experiments, where the sensitivity of different strains to toxic substances, environmental stresses such as temperature changes can be compared. Indeed, as the microsatellite barcode procedure can be easily replicated on batches of unique polyps, it represents a major asset for discriminating among phenotypically similar polyps those that are genetically different, and vice versa.

### Possible mechanisms explaining the strain-specific variations observed in Hydra microsatellite sequences

Karyotyping on *Hm-105* revealed that *Hydra* are diploid animals (2n=30) (55). It is therefore not surprising to observe either a single band or more frequently the same band completed by a second band, reflecting the homozygous versus heterozygous status of a given animal respectively. On the other hand, we interpret the differences in band size observed in animals of different strains as different alleles. Nevertheless, we have clearly observed and sequenced more than two different bands in the same polyp (see *ms-c25145 in Hm-105* and *Basel1*). As mentioned above, the *ms-c25145* primers we have designed can amplify two different regions of the *Hm-105* genome (*Sc4wPen_1246, Sc4wPen_396*), which explains why four bands can be observed in this strain (twice two alleles). The most parsimonious scenario would be that these two regions result from a recent single gene duplication that occurred in the common ancestor of the *Hm-105* and *Basel1* strains, without affecting the other strains tested here where only one copy is detected.

The microsatellite barcoding might also reveal some genetic mosaicism, as suspected from the four-band and three-band patterns observed for *ms-c25145*. Genetic mosaicism is defined as genetic variations acquired post-zygotically in cells of an individual developed from a single zygote, a phenomena frequently observed in plants and clonal animals as well as in humans (56,57). In clonal animals as cnidarians, the segregation of germ cells does not occur during early embryonic development and mutations affecting somatic cells as well as germ cells can accumulate over the multiple divisions of multipotent stem cells. In *Hydra*, beside the interstitial stem cell population that can transiently provide germ cells, the two epithelial stem cell populations also continuously cycle over the lifetime of the animal, potentially accumulating somatic mutations independently. This mechanism provides the opportunity for additional genetic variations within the same animal as observed in leaf cells (58).

### The microsatellite analysis supports the hypothesis of cryptic species within the H. vulgaris group

With the development of “omics” over the last decade, the subdivision of the genus *Hydra* into four main species, as initially proposed on morphological, developmental and cellular criteria, appears more complicated. Indeed, if molecular phylogenetic analyses have confirmed this four group classification, they have also highlighted the existence of several subgroups within each species (13–16), and one issue concerns the specification of separate species within the *H. vulgaris* group. For instance, Martinez et coll. (13) consider the *H. vulgaris* group as a single species clustering into five main sub-groups defined by their geographical distribution: South Africa, North America (e.g. *AEP*), South America, Eurasia (e.g. *Basel, Zürich, Hm-105*) and Oceania. Similarly, Kawaida et coll. (12) consider the *H. vulgaris* group as forming a single species but described three main sub-groups called *H. vulgaris*, *H. carnea* and *H. sp.*

In contrast, Schwentner and Bosch (14) suggest that the *H. vulgaris* group is more complex than expected, revealing at least 14 distinct subgroups, each representing a hypothetical species. They propose to cluster within a single *H. carnea* species the *H. carnea, H. littoralis* and the majority of the North American *H. vulgaris* strains, including the *AEP* strains. As a consequence, the *H. vulgaris* strains from Europe and Japan, named by these authors *H. vulgaris 1* would form a distinct *H. vulgaris* species, corresponding to that initially described in the 18e century by Trembley (1). The analysis of the microsatellite polymorphism reported in the present study supports this view as the data obtained on six wild-type strains that belong to the *H. vulgaris* group point to a divergence between the North American (*AEP*) and the Eurasian (*Hm-105, reg-16, Basel1, Basel2*) sequences. The acquisition of a genome for each sub-group would help to perform meta-analyses and analysis of single-nucleotide polymorphism to state on *H. vulgaris* species delimitation as recently done for the *Ophioderma* sea stars (59).

## CONCLUSION

With this study, we implemented a powerful barcoding approach based on microsatellite polymorphism for strains belonging to the *H. vulgaris* group. The use of this approach should enhance the reproducibility of experiments conducted in different laboratories by allowing the correct identification of each strain, including the AEP transgenic lines, in order to conduct unbiased experiments on well-characterized polyps. Data obtained on six wild-type strains belonging to the main hydra species used in experimental biology, namely *H. vulgaris* (referred to here as *H. vulgaris1*) and *H. carnea*, tend to confirm that the *H. vulgaris* group actually covers a set of cryptic species rather than a single species. We believe that microsatellite polymorphism analysis can help discover speciation events, thus representing a complementary approach to phylogenetic analyses aimed at identifying *Hydra* species.

## Acknowledgements

This work was supported by the Swiss National Science Foundation (SNF grants 31003_169930), the Claraz donation and the Canton of Geneva.

**Contributions**

QS proposed the initial concept and design of the project; QS identified the microsatellites and designed the primers. DPC and QS implemented the PCR approach and performed the PCRs. CP and QS prepared the samples for sequencing. BG supervised the research. BG and QS discussed the results and wrote the manuscript.

